# Immunogenicity of Rabies Virus G-Protein mRNA Formulated with Muscle Targeting Lipid Nanoparticles in Mice

**DOI:** 10.1101/2025.01.06.631441

**Authors:** Qin Li, Huarong Bai, Xueliang Yu, Qiang Liu, Rongkuan Hu

**Affiliations:** Starna Therapeutics Co., Ltd., Suzhou, 215123, China

**Keywords:** Rabies virus, mRNA vaccine, Muscle delivery, Lipid nanoparticles

## Abstract

Rabies is a preventable zoonotic disease caused by the rabies virus (RABV) with a high mortality rate. Most vaccines on the market or under development have issues such as a low single-dose neutralization titer, complex processes, and high costs. During the COVID-19 pandemic, the successful development of mRNA vaccines has opened up a new avenue for preventive vaccines. As a new technology, mRNA has higher scalability. In this study, we designed an mRNA encoding the RV-G protein, encapsulated by our own muscle targeting lipid nanoparticles (LNP), and evaluated the expression of the RV-G protein in vitro, its immunogenicity, and its protection against virus infection in vivo. The results showed that RV-G mRNA was significantly expressed in vitro. High Virus-IgG binding titers and Virus-neutralizing antibody titers (VNT) were induced by immunization with RV-G mRNA-LNP. Additionally, our results show that the RV-G mRNA vaccine is better than commercially available vaccines in mice.

## Introduction

Rabies is a fatal disease caused by the rabies virus (RABV). RABV invades the host’s central nervous system and causes pathological changes. Since the virus cannot be eliminated by the immune system or any drugs, once rabies occurs, the fatality rate can reach 100%.

RABV is a neurotropic virus of the genus Lyssavirus in the family Rhabdoviridae. It has a single-strand negative-sense RNA genome and five structural proteins: nucleocapsid protein (N), phosphoprotein (P), matrix protein (M), glycoprotein (G), and viral RNA polymerase (L) [1]. Among them, G protein is the only surface-exposed viral protein on RABV virions and it is the only virus protein that stimulates virus neutralizing antibodies[2].

The disease has been an important public health problem with no effective treatment to date. Prevention is mainly through vaccination currently. Inactive rabies vaccines are the most common and widely used around the world. They are produced in human diploid cells, chicken embryo cells, and Vero cells[3–6]. However, the cost of this vaccine is high. The main reason is due to the low titer of the rabies virus produced, which requires multiple injections to reach a higher level and titer so as to obtain satisfactory protective efficacy. Novel DNA vaccines, subunit vaccines, or viral vector vaccines have been evaluated to protect against rabies infection in preclinical settings, but these vaccines have not been approved for use in humans yet[7–9].

Messenger RNA (mRNA) vaccines are a revolutionary vaccine that has received a great deal of attention and use, especially during the COVID-19 pandemic, showing great promise and opening up new avenues for vaccine development and pandemic preparedness[10, 11].

In this study, we developed an mRNA -based rabies vaccine. This approach yielded high titers of virus-neutralizing antibodies and virus-IgG binding antibodies in mice, and it demonstrated effective protective efficacy in subsequent challenge experiments.

## Results

### Optimization of RV-G mRNA

Based on the key elements of mRNA (Figure 1A), we optimized the RV-G mRNA sequence. The specific optimization strategies are outlined in Table 1.

**Figure 1.**
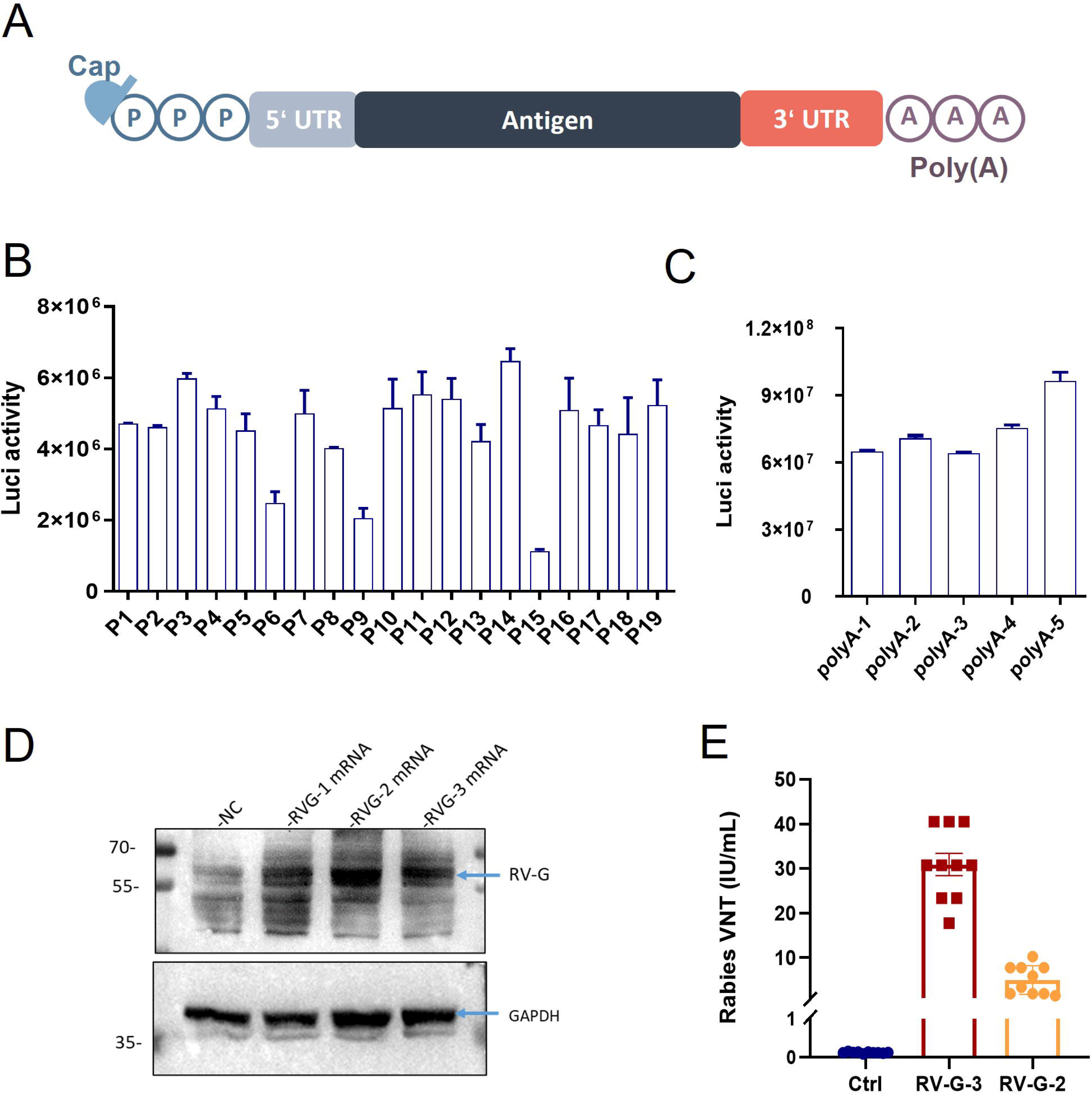
Optimization of RV-G mRNA sequence. A: key elements of synthetic mRNA; B: Screening results of UTR pairs based on the luciferase sequence; C: Results of PolyA optimization based on the luciferase sequence; D: Expression of RV-G protein for the three RV-G RNA variants; E: Virus-neutralizing antibody titers in the serum

**Table 1:**
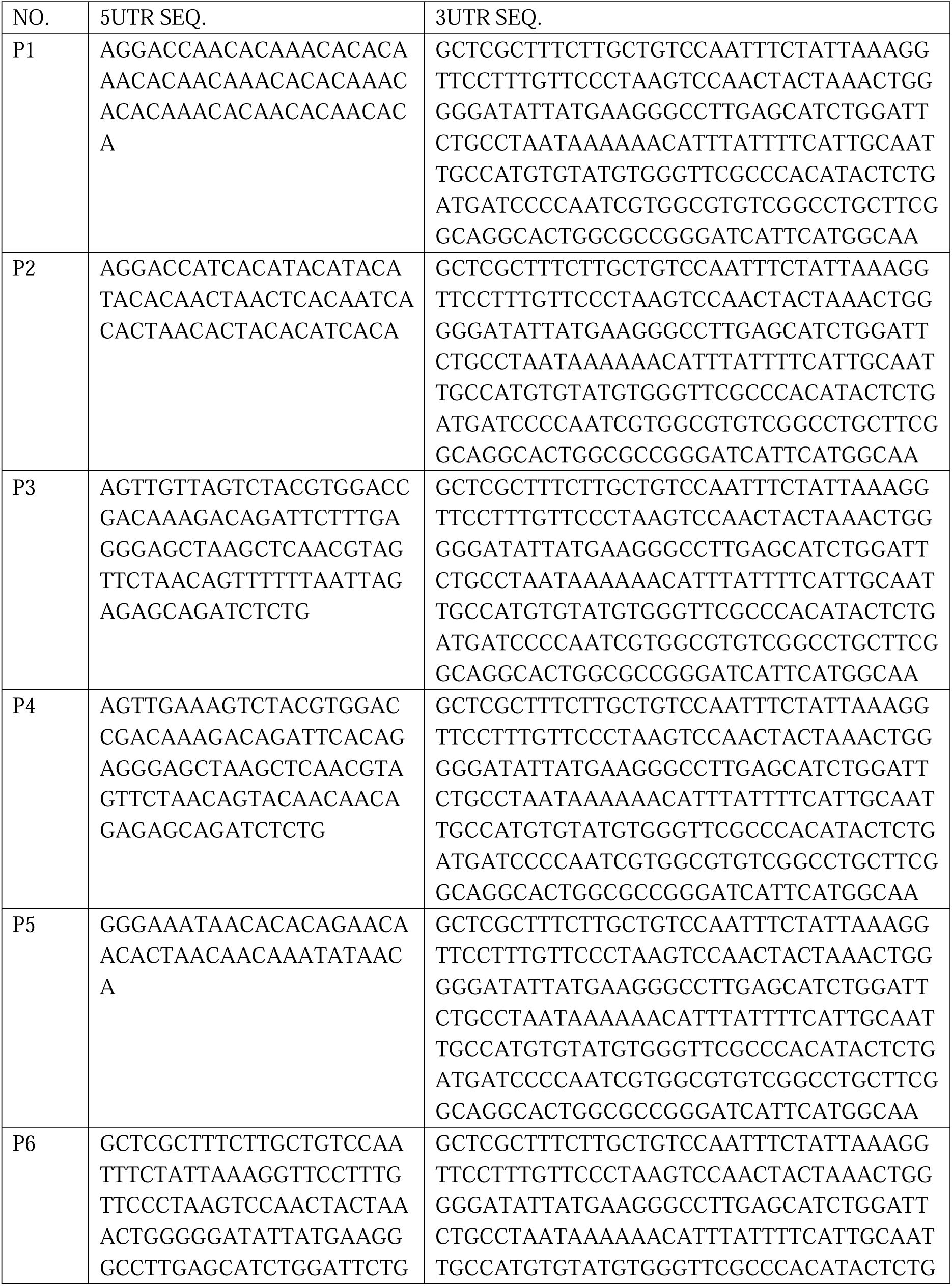
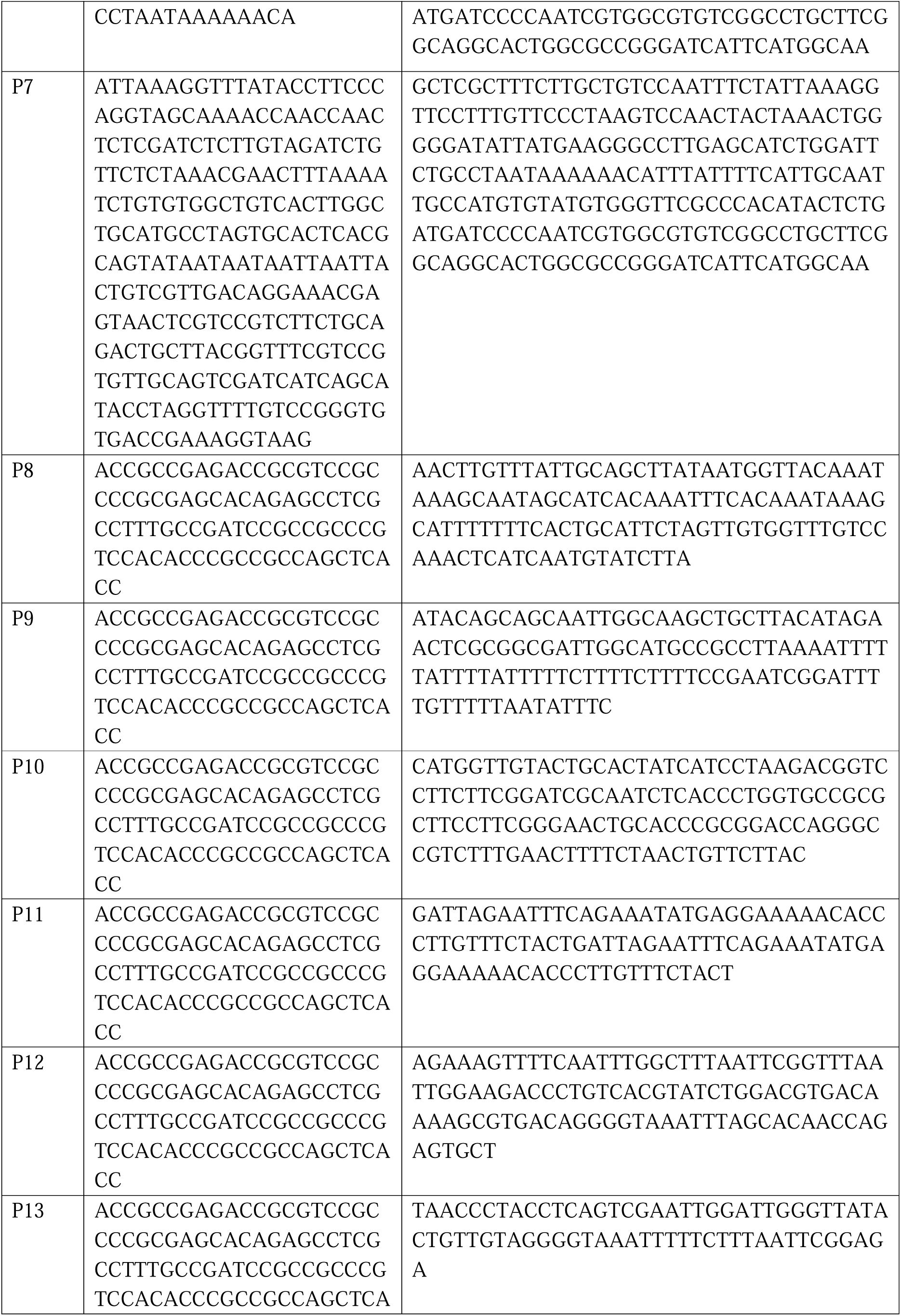
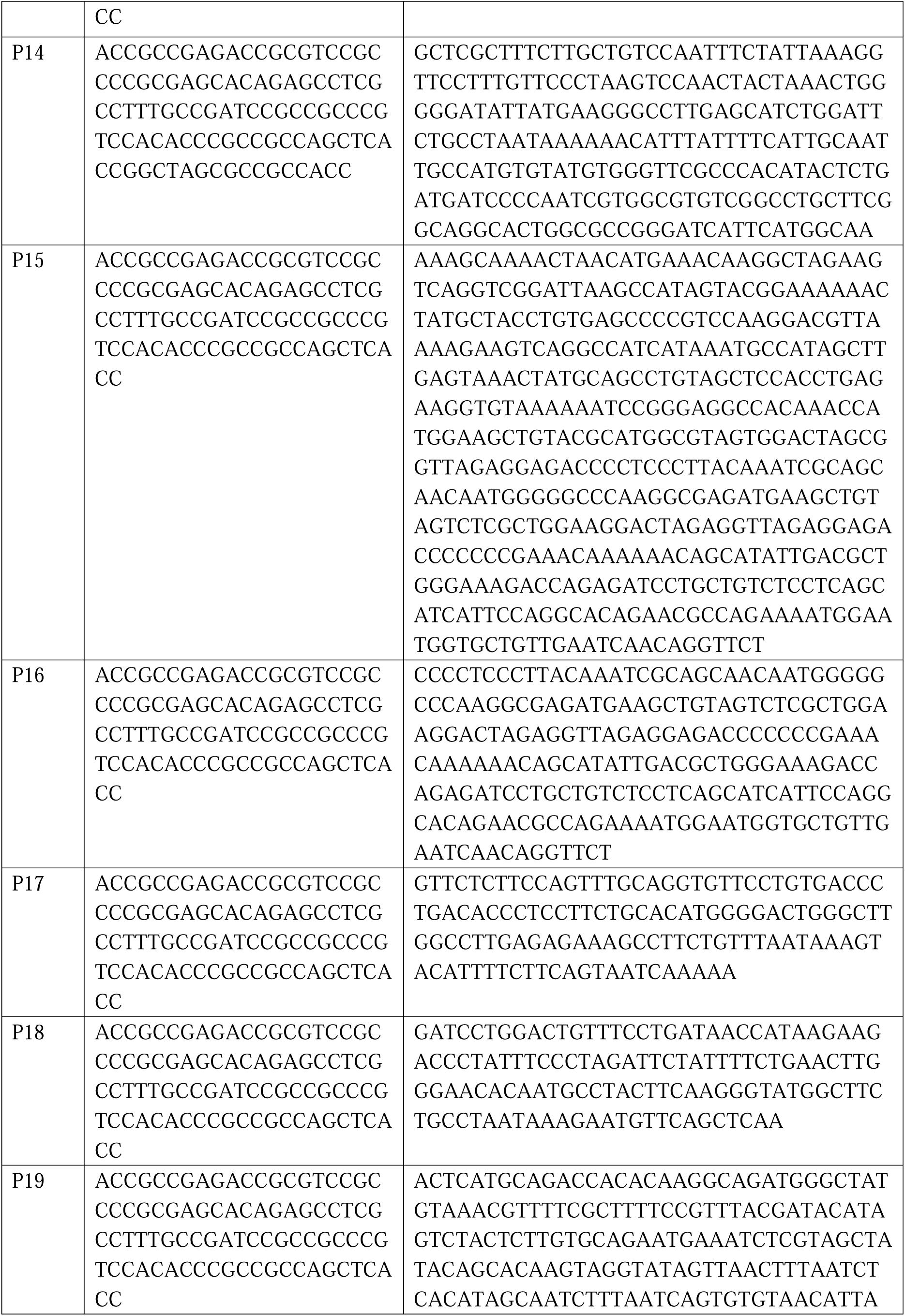
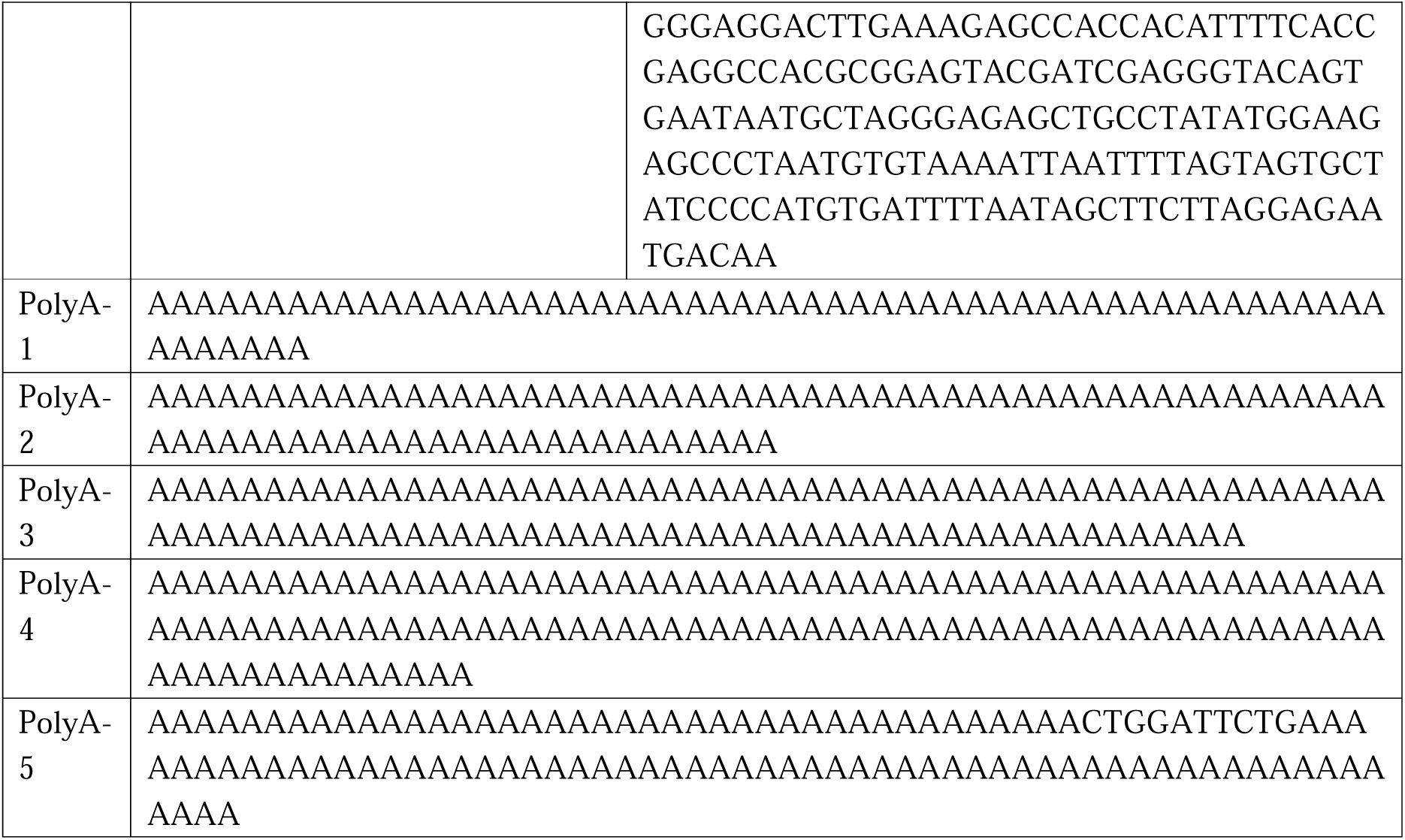
UTR sequences for UTR pair screening.

We initially screened for appropriate UTR pairs in the proprietary UTR library of Starna. These UTR pairs encompassed natural sequences, optimized sequences, AI-generated sequences, and de novo designed sequences. The complete list of UTR pairs can be found in Supplementary Information (Table 1). After selecting the UTR pairs, we employed the luciferase sequence as a coding region for initial screening, capitalizing on its straightforward manipulation. Different UTR pairs were incorporated into luciferase mRNA, which was then transfected into cells, and the luciferase (luc) activity was measured. The outcomes revealed excellent luciferase expression across all UTR pairs (Figure 1B), indicating that all UTR pairs were of highly effective when integrated into luciferase mRNA with P14 to be the best pair.

Subsequently, we screened the application of polyA. The candidates included full-length polyA and fragmented polyA. The specific sequences are shown in the Supplementary Information (Table 1). Results showed that the fragmented polyA showed a higher expression level (Figure 1C). So far, we have screened out the preferred UTR pair and the polyA.

Afterwards, we replaced the luciferase sequence with three optimized RV-G RNA sequence. These RV-G RNA variants were transfected into cells, and Western blot analysis was employed to detect RV-G expression. The findings indicated that the expression levels of RV-G in RV-G-1, RV-G-2, and RV-G-3 were markedly higher than that of the blank control in HEK-293T cells, confirming the successful expression of the RV-G protein by these RV-G RNA constructs (Figure. 1D). And in the in vivo experiments, RV-G-3 RNA induced neutralizing antibody titers were much higher than those provoked by RV-G-2 RNA after a single injection of 5ug (Figure. 1E). Therefore, we selected the RV-3 mRNA, which demonstrated better protein expression and higher neutralizing antibody titers, for further research.

### Muscle targeting LNP formulation screening

For the screening of muscle targeting LNP formulation, we introduced five distinct LNP formulations and evaluated the in vivo expression and safety-related immune responses. STAR-001, a traditional liver-targeting lipid nanoparticle (LNP), was used as a positive control, while for the other LNPs, cationic lipid was used as described previously [12, 13]. Mice were administered mRNA encoding luciferase (Luc) encapsulated within these varying LNP formulations via intramuscular injection. Six hours post-injection, luciferase expression in the mice was monitored by a bioluminescence imaging system. The findings revealed that all mice injected with Luc mRNA-LNP demonstrated luciferase expression, with the STAR-001 formulation showing the highest efficacy in liver (Figure 2A). In this research, we focused on developing mRNA vaccine formulations optimized for intramuscular injection rather than liver targeting, we hope to observe more enrichment at the site of muscle injection. As shown in figure 2B and table 2, STAR-002 formulation has the highest muscle to liver ratio.

**Figure 2.**
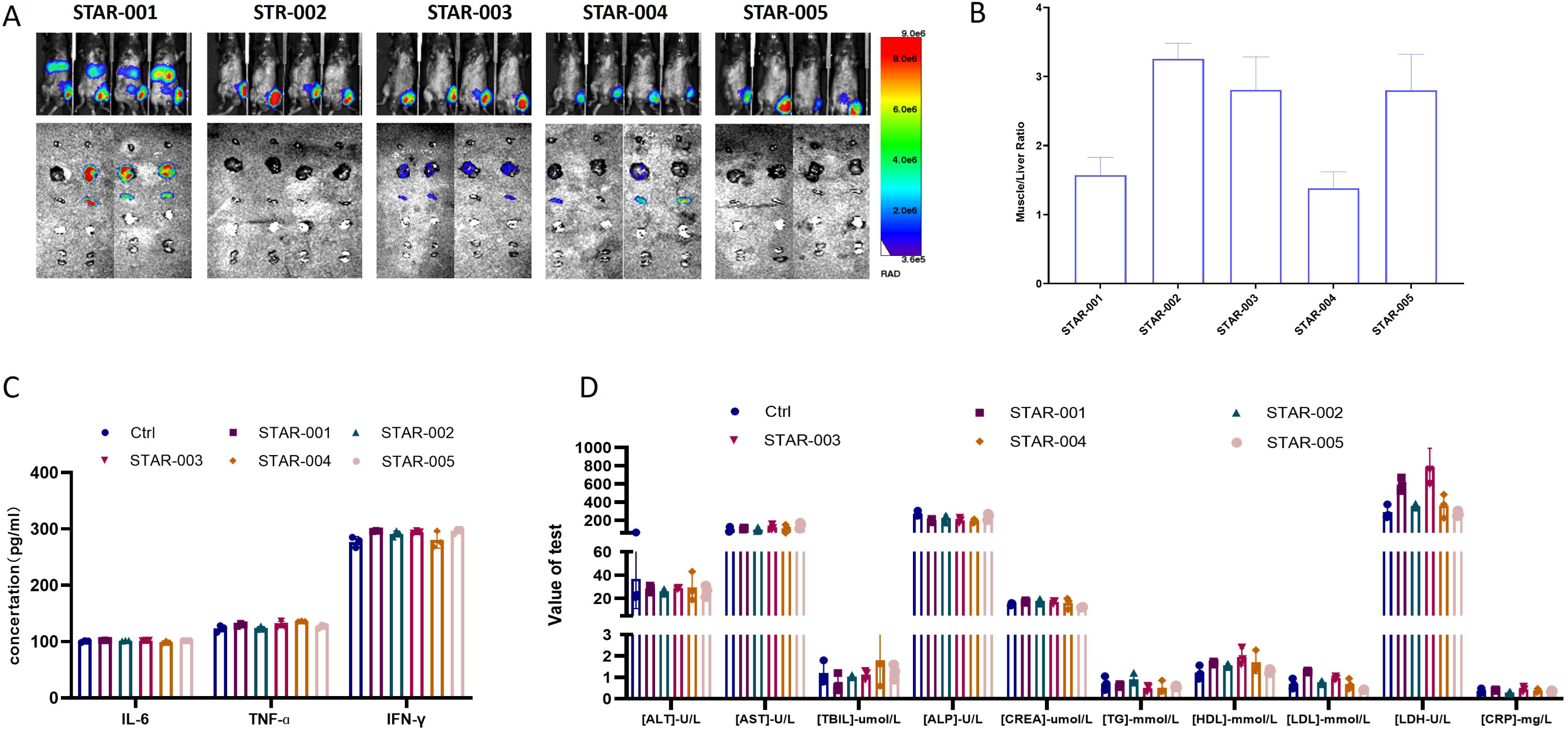
Muscle targeting LNP formulation screening. A: 6h after injection, luciferase expression was visualized in vivo; B: Muscle to liver ratio was calculated; C: Results of inflammatory cytokines assay; D: biochemical test results of the plasma.

**Table 2:**
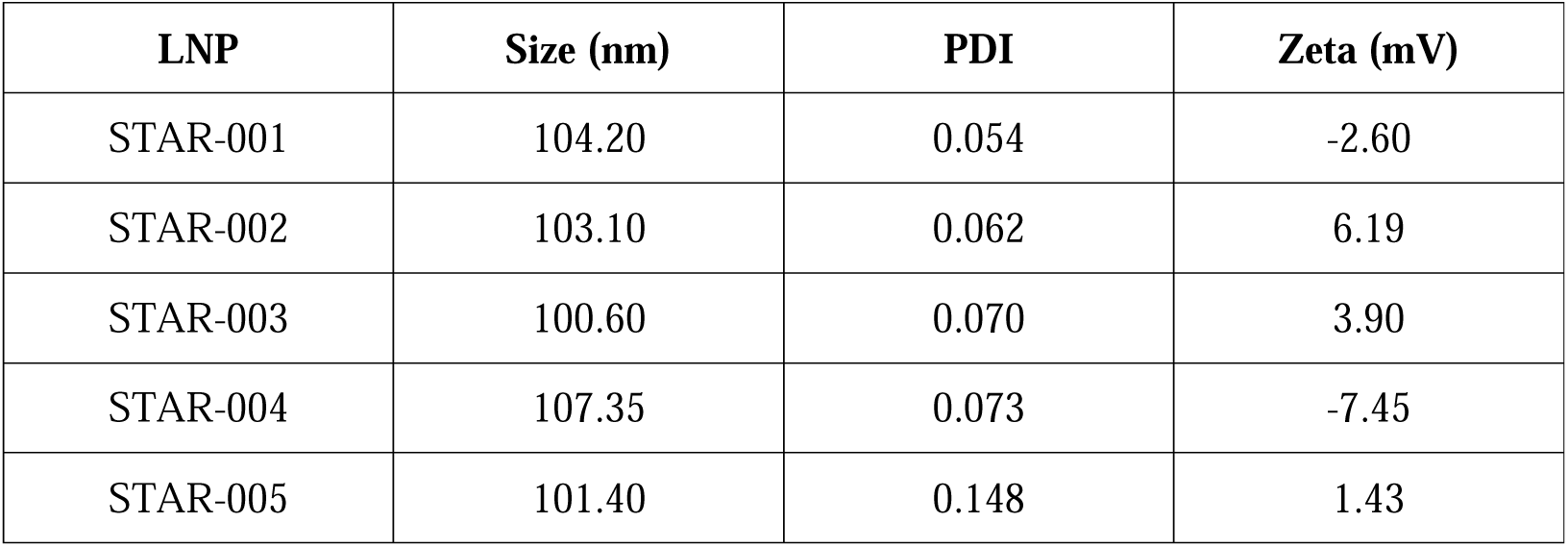
Size, PDI and zeta of LNP formulations.

To further assess the tolerability and safety of the LNP formulations, blood samples were collected one week after injection and analyzed for safety-related indicators. The data indicated that the serum concentrations of inflammatory cytokines (IL-6, TNF-α, and IFN-γ) and the plasma’s biochemical test results for most mice were similar to those in the mice injected with PBS (Figures 2C-D). Moreover, no significant weight loss was observed throughout this study. Collectively, these results suggest that all the LNP formulations mediate effective mRNA delivery and expression, and exhibit excellent tolerability and safety profiles. The STAR-002 formulation, exhibiting superior muscle targeting efficiency, was chosen for further development.

### RV-G mRNA-LNP induces strong humoral responses in mice

We have demonstrated that the optimized LNP formulations effectively mediate mRNA delivery and expression, along with good tolerability and safety. Additionally, through rigorous UTR pairs screening, we have identified the confirmed RV-G mRNA sequence. Here, we encapsulated the confirmed RV-G mRNA in the four superior formulations to obtain the optimal RV-G mRNA-LNP. BALB/c mice were immunized with these RV-G mRNA-LNPs on day 0, and blood samples were collected via the orbital venous plexus on day 7 and 14 for serological studies. Neutralization assay were performed, revealing that that the four LNP formulations induced significantly higher Virus-neutralizing antibody titers (VNT)on days 7 and 14 post-injection compared to the PBS group (Figure 2A). Notably, the STAR-002 formulation stood out with higher VNTs and less variability among groups. Consequently, we selected this formulation for further investigations.

Subsequently, we evaluated the in vivo immune response of the selected RV-G mRNA-LNP. BALB/c mice were immunized with 0.5, 5, and 50 µg doses of RV-G mRNA-LNP on day 0. Serological studies were conducted using blood samples collected via the orbital venous plexus on days 7 and 14 post-injection. ELISA was used to measure Virus-IgG binding titers, and Neutralization assay were performed. Results showed that on day 14, mice in the 5 µg RV-G mRNA-LNP group exhibited higher Virus-neutralizing antibody titers (VNT) and Virus-IgG binding titers (1:128000) than the control group (Figure 2B-C). And the VNT titers on day 7 ranged from 4.5 IU/mL in the 0.5 µg dose group to 26 IU/mL in the 50µg dose group. By day 14, these titers increased to range from 22.4 IU/mL in the 0.5 µg dose group to 71.9 IU/mL in the 50µg dose group, demonstrating a positive dose-response relationship (Figure 2D). These findings suggest that mRNA-LNP encapsulated with RV-G mRNA can elicit higher levels of Virus-IgG binding titers and Virus-neutralizing antibody titers (VNT) against the RV-G protein.

### The pre-exposure protection of RV-G mRNA-LNP and is capable of competing with licensed vaccines in mice

To assess the prophylactic capacity of RV-G mRNA-LNP in vivo, we conducted an experiment using 6–8-week-old Balb/C mice (n=10, with an equal number of males and females). The mice were intracerebrally infected with 100-fold LD50 CVS-24 strain 14 days after receiving a single 5 µg dose of immunization. A licensed vaccine served as a positive control.

The results were striking: all mice in the RV-G mRNA-LNP group survived, whereas 6 out of 10 mice in the positive control group succumbed This indicates a 100% protection rate for the RV-G mRNA-LNP group, compared to a 40% protection rate for the positive control group (Figures 4A-B). Additionally, we observed varying body weight changes among the groups. The RV-G mRNA-LNP group exhibited the least weight fluctuation, followed by the positive control group, which experienced a 10-day period of weight loss post-infection before gradually recovering. In contrast, the control groups continued to lose weight until they died (Figure 4C).

**Figure 3.**
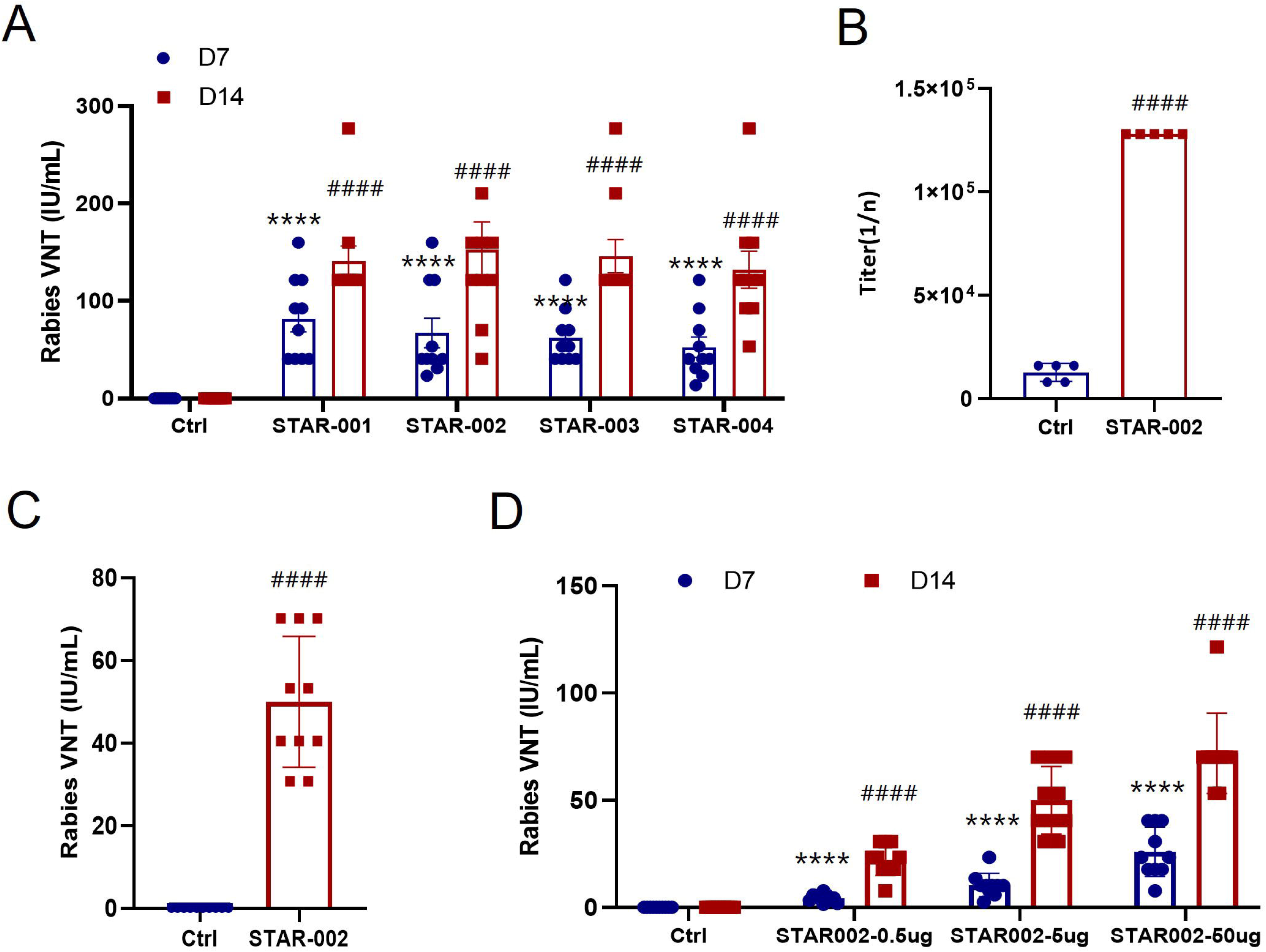
Antigen-specific antibody response in mice. A: Virus-neutralizing antibody titers in the serum of vaccinated mice detected by ELISA in the four superior formulations; B: Analysis of the antigen-specific antibody response in mice administered a 5 µg dose in serum collected on day 14; C: Virus-neutralizing antibody titers in the serum of the 5ug dose in serum collected on day 14; D: Virus-neutralizing antibody titers in the serum of the three dose in serum collected on day 7 and day 14on day 7 and day14 (****, P<0.001 compared to the Ctrl group on day7 ;####, P<0.001 compared to the Ctrl group on day14)

**Figure 4.**
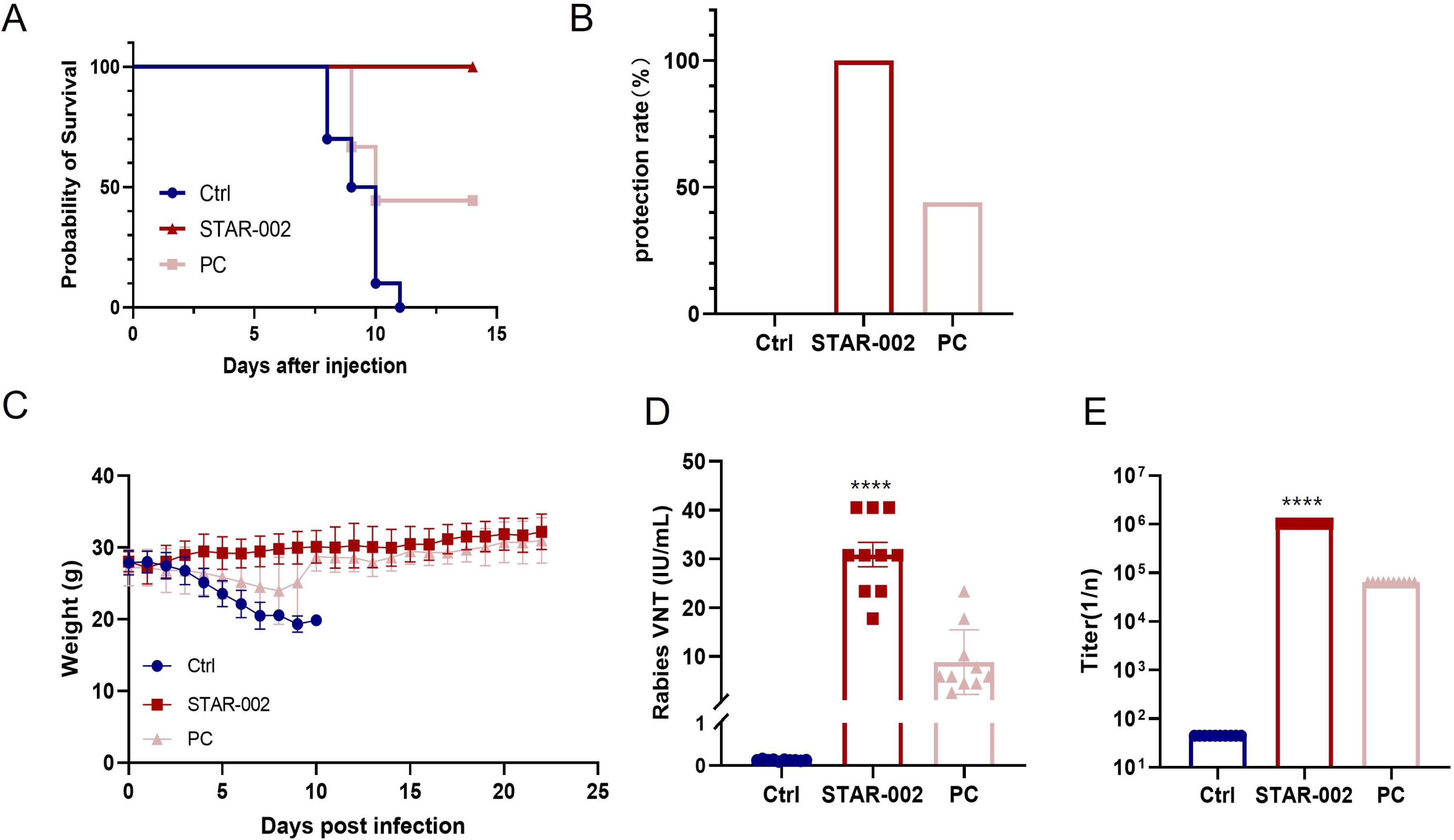
pre-exposure protection of RV-G mRNA vaccine in mice. A: Kaplan-Meier survival curves; B:Protection rate statistical chart; C: body weight changes in mice; D: Virus-neutralizing antibody titers in the serum; E: Virus-IgG binding titers in serum(****, P<0.001 compared to PC group)

We also compared the immunogenicity of RV-G mRNA-LNP to that of the licensed vaccine. A single 5 µg dose of RV-G mRNA-LNP induced Virus-neutralizing antibody titers (VNT) of 31 IU/ml, significantly higher than the 9 IU/ml induced by the licensed vaccine. Similarly, for Virus-IgG binding titers, the RV-G mRNA-LNP group showed a titer of 1:1024000, in contrast to the 1:64000 of the licensed vaccine (Figures 4D-E).

These findings suggest that a single 5 µg dose of RV-G mRNA-LNP results in Virus-neutralizing antibody titers (VNT) nearly 6 times the WHO standard of 0.5 IU/ml by day 14, which is sufficient to induce protective antibody titers.

### The post-exposure protection of RV-G mRNA-LNP

The aforementioned results clearly demonstrated that RV-G mRNA-LNP could protect mice from CVS-24 infection following pre-exposure immunization. Building on these findings, we sought to determine if RV-G mRNA-LNP also offers protection post-exposure. For this, we used 6–8-week-old Balb/C mice (n=10, with an equal number of males and females) and intramuscularly infected them with 50-fold LD50 CVS-24 6 hours prior to administering a single 5 µg dose of the immunization. Formulation SM-102 served as a positive control.

After infection, all groups experienced weight loss within the first 5 days; however, mice immunized with RV-G mRNA began to regain weight gradually, with the STAR-002 group demonstrating a more rapid recovery (Figure 5A). And the survival rates among the groups varied significantly. The STAR-002 group lost 4 mice, resulting in a 60% survival rate, while the SM-102 group lost 6 mice, yielding a 40% survival rate. In contrast, all mice in the control group succumbed to rabies within 14 days of infection (Figure 5B). Additionally, a single dose of STAR-002 and SM-102, both encapsulating RV-G mRNA, induced high levels of Virus-neutralizing antibody titers (VNT) 5 days post-administration, which were significantly higher than the WHO standard of 0.5 IU/mL (Figure 5C). These outcomes highlight that the STAR-002 formulation outperformed SM-102, providing superior protection against viral infection in vivo.

**Figure 5.**
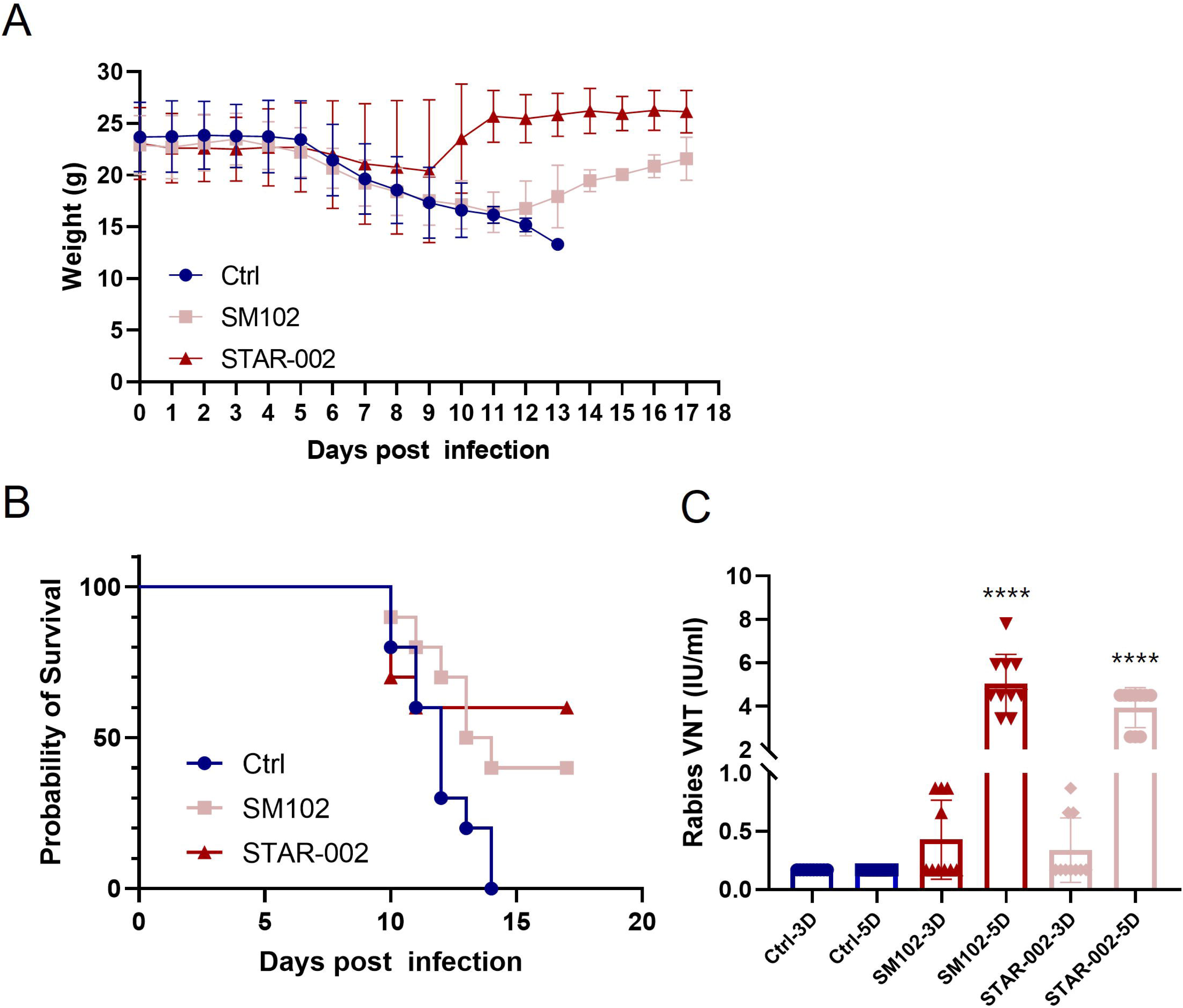
Post-exposure protection of RV-G mRNA vaccine in mice. A:body weight changes in mice; B:Kaplan-Meier survival curves; C:Virus-neutralizing antibody titers in the serum (****, P<0.001 compared to SM102 group)

## Discussion

mRNA represents a highly promising multifunctional vaccination approach, as underscored by its pivotal role in combating the COVID-19 pandemic. Two mRNA-based SARS-CoV-2 vaccines, mRNA-1273 (Moderna) and BNT162b2 (Pfizer-BioNTech), were swiftly developed, showcasing authentic safety and efficacy with relatively low costs[14, 15].

The inactivated rabies vaccine is prevalent globally, playing a crucial role in preventing and controlling the spread of rabies. However, most inactivated rabies vaccines necessitate multiple doses to achieve high neutralizing antibody titers, which incurs substantial costs. Thus, the development of innovative vaccines is desirable to help reduce the burden of rabies.

In this study, we engineered an RV-G mRNA vaccine focused on the rabies virus glycoprotein, the sole certified neutralizing antibody binding epitope. By optimizing the UTRs of RV-G mRNA, we enhanced the mRNA’s stability and efficiency. The RV-G mRNA, when encapsulated with various LNPs, induced high Virus-neutralizing antibody titers (VNT). Notably, the STAR-002 formulation provided 100% protection in pre-exposure challenge assays in mice, markedly outperforming the licensed vaccine’s 40% protection. In post-exposure challenge assays, it induced 60% protection, surpassing the classic formulation SM102, which offered 40% protection. Moreover, a single 5 µg dose of this vaccine yielded neutralizing antibody titers nearly 200 times the WHO standard of 0.5 IU/mL.

LNP-formulated mRNA can serve as a versatile platform technology[16], which can be manufactured efficiently through a standardized process. The simplicity and scalability of the LNP-formulated mRNA method, along with the ease of batch control, significantly reduce vaccine production costs. In conclusion, the encapsulation of RV-G mRNA as an antigen in LNP-based nanoparticles not only elicits Virus-neutralizing antibody titers (VNT) but also achieves significant immune effects with a single injection. These attributes render the mRNA vaccine a potentially safe and cost-effective candidate for rabies vaccination.

## Materials and Methods

### Cell lines

HEK293T cells, sourced from the Chinese Academy of Sciences Cell Bank in Beijing, China, and cos.7 cells, donated by Westlake University, were cultured in DMEM(HyClone), supplemented with 10%FBS(Gibco), and 1% Penicillin-Streptomycin (Gibco). Both of the cell lines were maintained under sterile conditions and incubated at a temperature of 37°C with a 5% CO2.

### mRNA design and preparation

UTR pairs were selected from the proprietary UTR library of starna. The coding sequence of the RV-G mRNA was obtained by performing a simple codon optimization on the RV-G sequence retrieved from GenBank (CTN-1 strain, Accession Number: AY009100.1). The construction of the DNA template for in vitro transcription was carried out by GeneScript (Nanjing, China). For mRNA preparation, the process began with in vitro synthesis using the T7 High Yield RNA Transcription Kit (Thermo, K0441). The resulting mRNA was then purified using the Monarch RNA Cleanup Kit (NEB, MA, UK).

### Lipid nanoparticle preparation

RV-G mRNA-LNP formulations were crafted using proprietary method. The lipid components, each with a defined molar ratio, were dissolved in ethanol, and the RNA was prepared in a 10 mM citrate buffer at a pH of 4.0. The synthesis process employed microfluidic technology, with an ethanol-to-water phase volume ratio set at 3:1 and a flow rate ratio of 1:3. Subsequently, a dialysis step was performed in 1×PBS for 2 hours to refine the formulation.

### mRNA transfection

HEK-293 cells were seeded in 96-well or 6-well plates one day prior to transfection. On the day of transfection, the cells were treated with 100 ng or 4 µg of luciferase or RV-G mRNA using Lipo2000 as the transfection reagent. After a 24-hour incubation period, the cells were harvested. Luciferase activity was measured by luciferase assay, and the expression of the RV-G protein was subsequently detected through Western blot and FACS analysis.

### luciferase assay

luciferase assay was carried out according to the Dual Luciferase Reporter Assay Kit (Promega E1910). Cells transfected with luciferase mRNA using Lipo2000 were harvested, the supernatant was removed, and the cells were rinsed once with 100 µl of PBS buffer, after which the buffer was discarded. Then, 105 µl of pre-diluted lysis buffer was applied to cover the cells and incubated at room temperature for 15 minutes. Subsequently, the 96-well plate was centrifuged at 13,000 rpm for 5 minutes. The supernatant was carefully transferred to a 96-well detection plate, with 30 µl aliquoted per well, and each sample was assayed in triplicate. Finally, 30 µl of luciferin substrate was added to each well in the dark and the luminescence values were read using a microplate reader (TECAN SPARK).

### Western-Blot

Cells transfected with RV-G mRNA using Lipo2000 were harvested, and the total protein concentration was measured using the Bradford assay. Following this, 1×SDS loading buffer was added to the samples, which were then subjected to SDS-PAGE and transferred onto PVDF membranes. The membranes were blocked with TBST containing 5% (w/v) nonfat dry milk for 2 hours at room temperature before being incubated with horseradish peroxidase (HRP)-conjugated secondary antibodies for 1 hour at 37°C. After three washes with 1X PBST, the blots were developed using a developing solution (Tanon, 180-5001). The resulting images were captured using a Tanon 4200 imaging system.

### Flow cytometric analysis (FACS)

After a 24-hour transfection with RV-G mRNA, cells were harvested by digestion with 0.25% trypsin. The cells were then re-suspended in 300μL of PBS and washed. This washing step was repeated twice with PBS. Subsequently, the Anti-Rabies Virus Glycoprotein antibody (Absolute Antibody, #Ab02097-3.0) was added to the cell suspension and incubated at 4°C for 2 hours. Following further washing with PBS, the cells were incubated with Fluor® 488 IgG H&L (Abcam, #ab150113) at 4°C for 1 hour. Flow cytometry (FACS) was used to analyze the cells. The flow cytometric data were quantitatively evaluated using FlowJo software.

### Mice immunizations procedure and Challenge infection

After a 3-day acclimation period, groups of six 6–8-week-old Balb/C mice were intramuscularly (i.m.) injected with 50 µL of different RV-G mRNA-LNP formulations, each containing 5 µg of mRNA. Plasma and serum samples were collected at 7-and 14-days post-injection for serological testing.

For the pre-exposure protection study of RV-G mRNA-LNP, mice were intracerebrally injected with 100-fold of the 50% lethal dose (LD50) of RABV CVS-24 strain 14 days after the immunization procedure. Serum was collected on day 14 for neutralization assays and to determine Virus-IgG binding titers.

In the post-exposure protection study of RV-G mRNA-LNP, mice were intracerebrally injected with 100-fold of the 50% lethal dose (LD50) of RABV CVS-24 strain 6 hours prior to the immunization procedure on day 0. Serum was collected on day 14 for neutralization assays. Following the challenge infection, the survival rates and body weight changes of the mice were monitored daily.

### Inflammation cytokines and biochemistry test

Serum samples were analyzed for inflammation biomarkers (IFNγ, IL-6 and TNF-LJ) using the Elisa kit (abcam # ab282874, #ab222503, # ab208348). Plasma biochemical test were carried out using automatic biochemistry analyzer (HITACHI 7100+ISE)

### Enzyme-linked immunosorbent assay (ELISA)

IgG titers in the immunized sera were assessed using an ELISA method. ELISA plates were coated with recombinant Anti-Rabies Virus antibody (final concentration: 2 µg/mL) (Abcam, #ab128726) and incubated at 4°C overnight. After washing with PBST twice, the plates were incubated with diluted serum following a blocking step with PBS containing 2% BSA for 2 hours at 37°C. Subsequently, the plates were incubated for an additional 2 hours at 37°C with an anti-Rabies Virus Glycoprotein conjugated to HRP (Abcam, #ab193430) to measure the specific binding of antibodies to the RV-G protein using a TMB substrate.

### Neutralization assay

Rabies virus challenge virus standard strain (CVS-11) was propagated on BSR cells and utilized in the experiments, following the Reed-Muench method. The standard serum was diluted to a concentration of 0.5 IU/mL with DMEM. A volume of 100 μL DMEM medium was added to each well of a 96-well plate, followed by the addition of 50 μL of the tested serum, standard serum, or negative control serum to the first column. Subsequently, a 3-fold gradient dilution was carried out across the plate. The test serum was diluted through column 9, while the standard and negative sera were diluted through column 6, with 50 μL being discarded from the final column. The CVS-11 virus was diluted to a concentration of 100 FFU per 50 μL with DMEM, and 50 μL of this virus dilution was added to each well containing the sera. Then, 100 μL of a BSR cell suspension containing 2×10^4 cells were added to each well and incubated in a 5% CO2 incubator at 37°C for 60 hours.

The culture medium was then removed from the 96-well plate, and the wells were fixed with precooled acetone at -20°C for 30 minutes and washed three times with PBS. The plate was incubated with a FITC-conjugated antibody specific for the rabies virus glycoprotein (RV-G) at 37°C for 45 minutes. After discarding the antibody incubation solution and washing three times with PBS, the cells were observed under a fluorescence microscope. The absence of fluorescence was recorded as “-”, while the presence of one or more fluorescent cells was recorded as “+”. The neutralizing antibody titers were determined accordingly. The Virus-neutralizing antibody titers (VNT) were calculated using the Reed-Muench method.

### Data analysis

Data analysis was conducted using GraphPad Prism version 8.0.2 for Windows (GraphPad Software, Inc., La Jolla, CA, USA). t-tests were employed to assess statistical significance. Kaplan-Meier survival curves were utilized to evaluate the statistical significance of survival rates post-challenge. The thresholds for statistical significance are denoted as follows: * for p < 0.05, ** for p < 0.01, *** for p < 0.001, and **** for p < 0.0001; “ns” indicates no significant difference.

## Acknowledgements

We thank employees of Starna Therapeutics for their helpful technical and scientific support with materials preparation. This study was supported in part from Starna Therapeutics (STR-V003), Jiangsu Provincial Science and Technology Project (SBK2023070025), Suzhou Science and Technology Project (ZXL2023267), the “Open Competition to Select the Best Candidates” Key Technology Program for Cell and Gene Therapy of NCTIB (NCTIB2023XB02001).

## Author contributions

R.H., and Q.L. conceived the project and designed the experiments and wrote the manuscript. R.H. and X.Y. supervised the project. H.B. and Q.L. carried out and participate in the whole molecular experiments.

